# A Thousand Fly Genomes: An Expanded *Drosophila* Genome Nexus

**DOI:** 10.1101/063537

**Authors:** Justin B. Lack, Jeremy D. Lange, Alison D. Tang, Russell B. Corbett-Detig, John E. Pool

**Author notes:** Corresponding author: John E. Pool, 425-G Henry Mall, Madison, WI 53706. Current address: Center for Cancer Research, National Cancer Institute, NIH, Bethesda, MD 20892-1201. Department of Biomolecular Engineering, University of California, Santa Cruz, Santa Cruz, CA 95604.

## Abstract

The *Drosophila* Genome Nexus is a population genomic resource that provides *D. melanogaster* genomes from multiple sources. To facilitate comparisons across data sets, genomes are aligned using a common reference alignment pipeline which involves two rounds of mapping. Regions of residual heterozygosity, identity-by-descent, and recent population admixture are annotated to enable data filtering based on the user’s needs. Here, we present a significant expansion of the *Drosophila* Genome Nexus, which brings the current data object to a total of 1,122 wild-derived genomes. New additions include 306 previously unpublished genomes from inbred lines representing six population samples in Egypt, Ethiopia, France, and South Africa, along with another 193 genomes added from recently-published data sets. We also provide an aligned *D. simulans* genome to facilitate divergence comparisons. This improved resource will broaden the range of population genomic questions that can addressed from multi-population allele frequencies and haplotypes in this model species. The larger set of genomes will also enhance the discovery of functionally relevant natural variation that exists within and between populations.

---

The genetics model *Drosophila melanogaster* has played a pivotal role in population genetic research. A growing number of studies have generated population genomic data from this species, but alignment and filtering criteria typically vary among studies, which obscures direct comparisons between these data sets. The *Drosophila* Genome Nexus (DGN; Lack *et al.* 2015; http://www.johnpool.net/genomes.html) provides the research community with genomes from multiple published sources that are generated using a common reference alignment pipeline. This pipeline improved upon typical reference alignment protocols by including a second round of mapping to a modified reference genome that incorporates the variants detected in the first round, yielding improved genomic coverage and accuracy (Lack *et al.* 2015).

Version 1.0 of the DGN included 623 genomes of *D. melanogaster* from individual wild-derived strains, originating primarily from three data sets. Phase 2 of the *Drosophila* Population Genomics Project (DPGP; Pool *et al.* 2012) included 139 genomes from 22 populations, mainly from Africa. *D. melanogaster* was known to have originated in sub-Saharan Africa (Lachaise *et al.* 1988), and this study identified southern-central Africa as the likely ancestral range. It also identified significant recent gene flow re-entering Africa, potentially related to urban adaptation, and powerful effects of inversions on genomic variation (Pool *et al.* 2012). Phase 3 of DPGP focused on a putative ancestral range population identified in the previous study, and brought this Zambia sample to a total of 197 independent, haploid genomes from a single location (Lack *et al.* 2015). That study, which also introduced the DGN, published additional African genomes from other populations, and confirmed that the focal Zambia sample was maximally diverse among all sampled populations, with minimal presence of non-African admixture (Lack *et al.* 2015). The third main data source of DGN 1.0 was from the *Drosophila* Genetic Reference Panel (DGRP), which consists of 205 genomes originating from Raleigh, North Carolina, USA (Mackay *et al.* 2012; Huang *et al.* 2014). These genomes were from strains inbred for 20 generations, resulting in 87% homozygous regions across euchromatic chromosome arms (Lack *et al.* 2015). North American populations appear to have resulted from admixture between European and African gene pools; a recent study that examined population ancestry along DGRP genomes estimated this population to be 20% African, with significant genome-wide evidence for incompatibilities between African and European alleles at unlinked loci (Pool 2015). Beyond these three main sources, DGN 1.0 also included Malawi genomes from DPGP Phase 1 (Langley *et al.* 2012) and source strain genomes from the Drosophila Synthetic Population Resource (DSPR; King *et al.* 2012).

In the present release, labeled as version 1.1 of the DGN, we add a total of 499 genomes. Of these, 306 are newly published in this study, and were sequenced from strains inbred for eight generations. These genomes were added to much smaller samples of genomes originating from a pair of Ethiopian populations (EA, EF), a pair of South African populations (SD, SP), and populations from Egypt (EG) and France (FR). These genomes facilitated a population genomic analysis of parallel adaptation to cold environments in three geographic regions, as described in an accompanying article (Pool *et al.* 2016). Genomic sequencing was performed using identical methods to those described by (Lack *et al.* 2015). Briefly, for each inbred line, ~30 female flies were used to prepare genomic DNA libraries. Sequencing on a HiSeq 2000 was performed to generate paired end 100 bp paired end reads with ~300 bp inserts.

DGN 1.1 also adds 193 genomes from four published studies. The Global Diversity Lines (Grenier *et al.* 2015) include 85 genomes from Australia, China, the Netherlands, the USA, and Zimbabwe. The 50 genomes published by Bergman and Haddrill (2015) originate from France, Ghana, and the USA. Campo *et al.* (2013) studied 35 genomes from a California population. Kao *et al.* (2015) added 23 genomes originating from 12 New World locations.

The population samples represented in DGN1.1 are depicted in Figure 1 and described in Table S1. Characteristics of all 1,122 individual strain genomes are given in Table S2. Instead of just three geographic population samples with more than ten sequenced genomes (as in DGN 1.0), there are now a dozen such samples (Figure 1), with five of these having more than 60 genomes.

**Figure 1.**
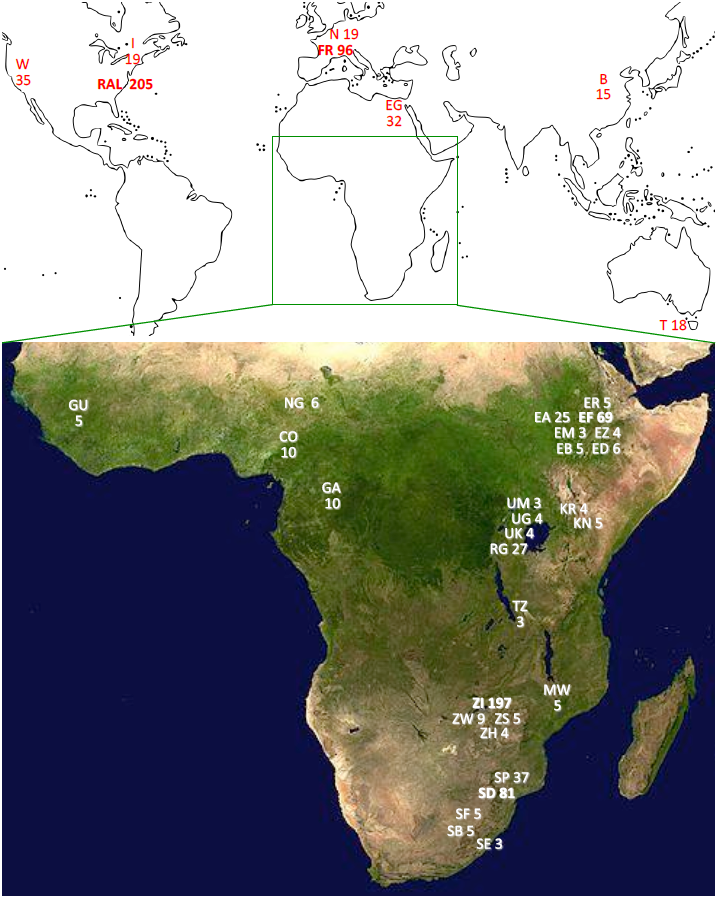
Geographic locations of sequenced population samples are shown, with the largest samples in bold print. These populations have at least three sequenced genomes with DGN consensus sequences available.

Importantly, the current DGN release does not modify the genomes represented in the prior data object. Instead, we have aligned and filtered the newly-added genomes using exactly the same pipeline described for DGN 1.0, again using the Flybase release 5.57 *D. melanogaster* reference genome. (Lack *et al.* 2015). Beginning with raw sequence read data, mapping is performed using BWA v0.5.9 (Li and Durbin 2010) followed by Stampy v1.0.20 (Lunter and Goodson 2010). GATK (Depristo et al. 2011) is then used to realign indels and generate consensus sequences. Called SNPs and indels are then incorporated into a genome-specific modified reference sequence, and read mapping is performed a second time to reduce mismatches. Genomic coordinates are then shifted back to match the original reference numbering. The “site” and “indel” variant call files (VCFs) provided by DGN are the direct output of this pipeline.

DGN also distributes consensus sequence files that feature additional filtering, and may be more appropriate for most analyses. To reduce the error rate, sites within 3 bp of a called indel are masked to “N”. For genomes that may contain residual heterozygosity, genomic intervals of apparent heterozygosity are fully masked. For fully haploid genomes (Langley *et al.* 2011), sites with an excess of apparent heterozygosity (e.g. due to technical artefacts or structural variation) are similarly masked as “pseudoheterozygosity”. Following such masking (in addition to removal of non-target chromosome arms from samples such as chromosome extraction line genomes), we find that an average site has homozygous consensus sequence calls from 754 DGN genomes.

We also provide files to enable user-initiated masking for two additional criteria. First, we allow regions of “identity by descent” due to relatedness between genomes in the same population sample to be masked. Second, we allow users to mask from sub-Saharan genomes regions of recent admixture from non-African populations (Pool *et al.* 2012). Full details on the alignment and filtering processes are given by Lack *et al.* (2015). Detailed filtering outcomes for heterozygosity, relatedness IBD, and admixture are provided in Table S3, Table S4, and Table S5, respectively.

Filtering characteristics of several data sets are depicted in Figure 2. Substantial heterozygosity persists in genomes sequenced from inbred lines (GDL, Campo, Kao, Pool, DGRP), in spite of inbreeding efforts that would be expected to reduce heterozygosity to nominal levels under neutral assumptions. Note that in Figure 2, “heterozygosity” also includes regions masked due to elevated heterozygous site rates for reasons such as copy number variation or data quality (“psuedoheterozygosity”; Lack *et al.* 2015). For example, the DGRP data set is estimated to have just 13% genuine heterozygosity (Lack *et al.* 2015). Previous analysis has shown that most genuine residual heterozygosity is associated with inversions (Grenier *et al.* 2015; Lack *et al.* 2015). Inversion genotypes based on prior published calls and the method of Corbett-Detig *et al.* (2012) are given in Table S6. Genomes from the Bergman and Haddrill (2012) data set, which were sequenced from isofemale lines, were estimated to be 99% heterozygous. DGN provides VCFs but not heterozygosity-filtered consensus sequences for these genomes.

**Figure 2.**
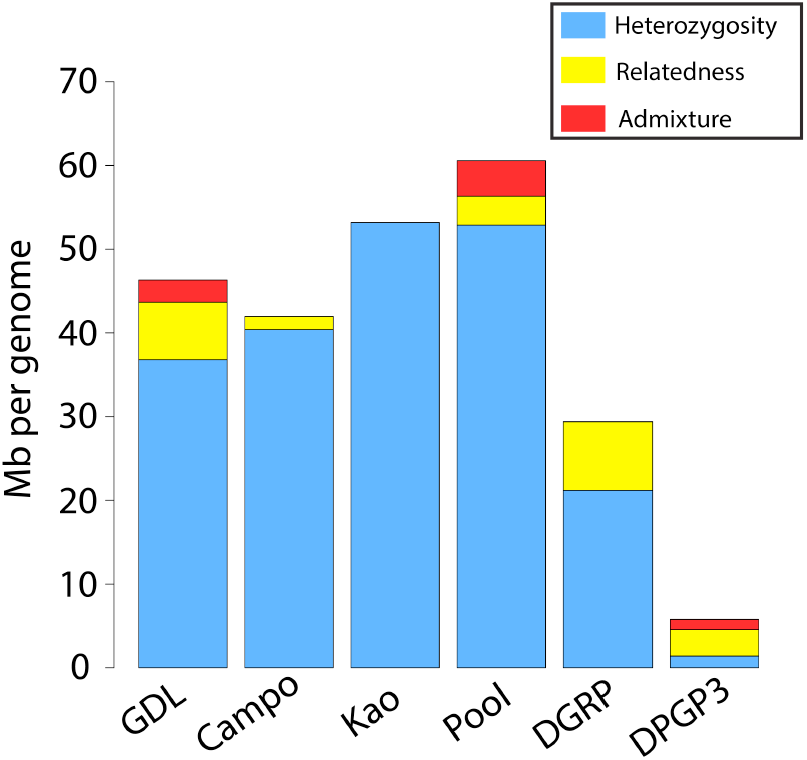
The extent of genomic data annotated for masking due to heterozygosity, relatedness, and admixture is shown per 119 Mb genome (when filtered in that order).

Figure 2 also shows the proportion of data sets that can be masked for relatedness IBD. IBD levels vary among population samples, with very high levels observed for the Netherlands GDL sample (where 30% of data would be masked for IBD, even though we only mask one member of each IBD pair), along with somewhat high IBD levels observed for the DGRP.

Pool *et al.* (2012) found evidence for substantial recent gene flow from non-African populations back into sub-Saharan genomes. Masking admixed genomic regions may allow sub-Saharan genetic diversity to be studied more directly, with fewer departures from typical assumptions of well-mixed populations. Admixture levels are known to vary drastically between sub-Saharan populations, partly as a function of urbanization (Pool *et al.* 2012). Of the data sets shown in Figure 2, “Pool” is mostly comprised of sub-Saharan genomes (62% from Ethiopia or South Africa), while one sixth of “GDL” consists of Zimbabwe genomes. “DPGP3” is a sample of 197 genomes from a single Zambia population with very low levels of admixture (Lack *et al.* 2015).

Among the DGN 1.1 samples, 13 worldwide populations are represented by at least 10 genomes for all three euchromatic chromosomes. A summary of genetic variation within and between populations is provided in Figure 3. As previously indicated, genomic diversity is highest in Zambia and other southern African populations (Pool *et al.* 2012; Lack *et al.* 2015), and all sub-Saharan populations are more diverse than all others. Because North American populations have mainly European but partly African ancestry (Kao *et al.* 2015; Pool 2015; Bergland *et al.* 2016), they show somewhat higher diversity than European populations. Geographic structure is apparent, especially between sub-Saharan populations and all others, with the latter group showing a common reduced gene pool apparently resulting from a population bottleneck. Additional bottlenecks may have impacted the B population from China (Laurent *et al.* 2011) and the EF population from the Ethiopian highlands (Pool *et al.* 2012; Lack *et al.* 2015), leading to mild population-specific reductions in diversity and increases in genetic differentiation (Figure 3).

**Figure 3.**
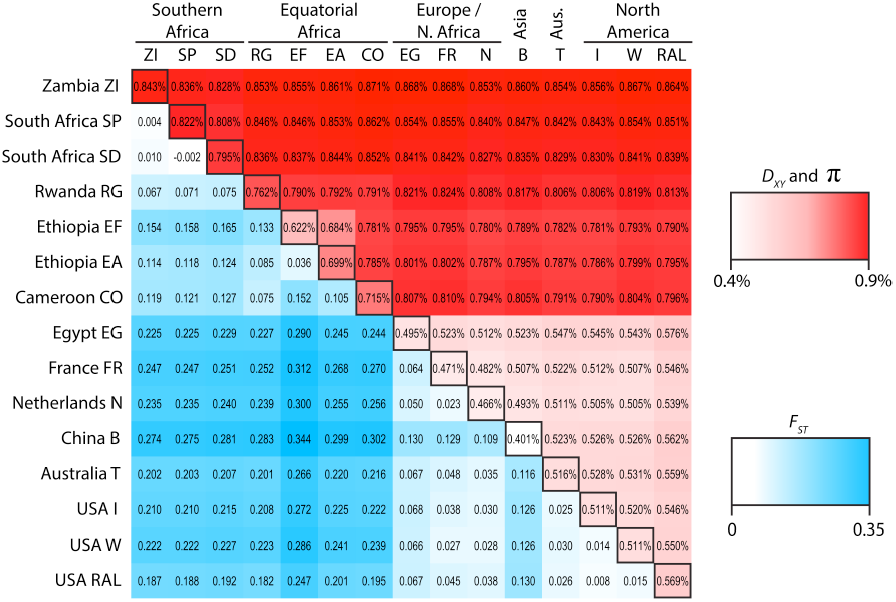
Average values of nucleotide diversity (*π*) within populations (on the diagonal), average pairwise distance between populations (*D*_*xy*_, above the diagnonal), and *F*_*st*_ between populations (below the diagonal) are shown. Values are averaged across chromosome arms X, 2L, 2R, 3L, and 3R, each of which was analyzed using inversion-free genomes only.

In addition to the above-described *D. melanogaster* genomes, DGN now also distributes an aligned sequence of *D. simulans* to the same *D. melanogaster* reference genome. Stanley and Kulathinal (2016) produced this alignment using progressiveMauve (Darling *et al.* 2010) to align the release 2 *D. simulans* genome (Hu *et al.* 2013) to the release 5 *D. melanogaster* reference sequence. We provide sequence text files mirroring our *D. melanogaster* consensus sequences for *D. simulans* on the DGN web site (http://www.johnpool.net/genomes.html). Note that for all data hosted by DGN, users should cite the original publications (Table S2) in addition to this alignment resource.

This expansion of the DGN will significantly bolster researchers’ ability to examine genetic variation within and between *D. melanogaster* populations. Future DGN releases will entail realigning all genomes using updated methods and reference genomes, plus evaluating new formats for providing genomic data. Community input to shape the future of this population genomic resource is welcome.

## ACKNOWLEDGMENTS

The UW-Madison Center for High Throughput Computing provided computational assistance and resources for this work. This research was funded by NIH grants R01 GM111797 to JEP and F32 GM106594 to JBL.

